# Impairment of brain function in a mouse model of Alzheimer’s disease during the pre-depositing phase: the role of α7 nicotinic acetylcholine receptors

**DOI:** 10.1101/2024.06.06.597737

**Authors:** Olena Lykhmus, Wen-Yu Tzeng, Lyudmyla Koval, Kateryna Uspenska, Elizabeta Zirdum, Olena Kalashnyk, Olga Garaschuk, Maryna Skok

**Author notes:** Corresponding author: Olga Garaschuk, Institute of Physiology, Department of Neurophysiology, University of Tübingen, Keplerstr. 15, 72074 Tübingen, Germany. These authors contributed equally to this work (Olena Lykhmus, Wen-Yu Tzeng). Deceased, April 11, 2024 (Maryna Skok).

## Abstract

Alzheimer’s disease (AD) is an age-dependent incurable neurodegenerative disorder accompanied by neuroinflammation, amyloid accumulation and memory impairment. It begins decades before the first clinical symptoms appear, and identifying early biomarkers is key for developing disease-modifying therapies. We show now in a mouse model of AD that before any amyloid deposition the brains of 1.5-month-old mice contain increased levels of pro-inflammatory cytokines IL-1β and IL-6, decreased levels of nicotinic acetylcholine receptors (nAChRs) in the brain and brain mitochondria and increased amounts of α7 nAChR-bound Aβ_1-42_, along with impaired episodic memory and increased risk of apoptosis. Both acute (1-week-long) and chronic (4-month-long) treatments with α7-selective agonist PNU282987, starting at 1.5 months of age, were well tolerated. The acute treatment did not affect the levels of soluble Aβ_1-42_ but consistently upregulated the α7 nAChR expression, decreased the level of α7- Aβ_1-42_ complexes and improved episodic memory of 1.5-month-old mice. The chronic treatment, covering the disease development phase, strongly upregulated the expression of all abundant brain nAChRs, reduced both free and α7-coupled Aβ_1-42_ within the brain, had anti-inflammatory and antiapoptotic effects, and potently upregulated cognition, thus identifying α7 nAChRs as both early biomarker and potent therapeutic target for fighting this devastating disease.

## 1. Introduction

Alzheimer’s disease (AD) is an incurable neurodegenerative disorder and the main cause of dementia in the elderly. According to the most recent data from the World Health Organization, the number of people living with dementia was estimated at 55 million in 2019 and was expected to rise to 139 million in 2050 (https://www.alzint.org/resource/world-alzheimer-report-2023/). These estimations, however, were made before the COVID-19 pandemic. The latter has infected more than 540 million people worldwide and the typical features of COVID-19, including hypoxia, cytokine storm, systemic inflammation and damage of the blood-brain barrier often result in delirium, brain fog and cognitive dysfunction even in young patients without comorbidities, thus further increasing the risk of developing AD [1–3].

Amyloid plaques represent the main histological hallmark of the disease, and many therapeutic approaches are targeted against amyloid β (Aβ) accumulation and oligomerization [4, 5]. Besides amyloidosis, AD patients present with an accumulation of the neurofibrillary tangles formed by the hyperphosphorylated microtubular tau protein, neuroinflammation and neurodegeneration [6–8]. Moreover, neurons iPSC-derived from AD patients show dysregulated intracellular Ca^2+^ homeostasis [9], in line with data obtained in various AD mouse models [10–15].

Another line of evidence relates the development of AD pathology to cholinergic transmission. Indeed, the source of cortical cholinergic innervation in the basal forebrain degenerates early during the disease progression [16] and cholinesterase inhibitors, increasing the availability of acetylcholine at brain synapses, represent one of the few clinical AD therapies. The nicotinic acetylcholine receptors (nAChRs) are ligand-gated ion channels mediating fast synaptic transmission in the neuro-muscular junctions and autonomic ganglia [17]. They are also expressed in the brain, where they regulate neurotransmitter and cytokine release [18]. Structurally, the neuronal nAChRs are homo- or heteropentamers composed of either identical α subunits (α7 or α9) or combinations of α2-α10 and β2, β4 subunits. The subtypes abundantly expressed in the brain and mostly related to AD are α4β2, known to control learning and memory [19], and α7 or α7β2 involved in APP processing, learning, memory and inflammation [20–22]. The α7 nAChRs are expressed in many CNS cells, including neurons, microglia, astrocytes and endothelial cells [23, 24]. They have a high Ca^2+^ permeability (8-12% of the current through the channel is carried by Ca^2+^), comparable to that of NMDA receptor channels [25], thus being able to activate several elements of second messenger pathways (e.g. protein kinases, NO synthetases, calcineurins, and other enzymes). Besides, α7 nAChRs can activate the JAK2-STAT3 signaling pathway thus inducing anti-apoptotic and anti-inflammatory effects as well as the AMPK-mTOR signaling pathway to induce autophagy [24]. They also mediate the cholinergic anti-inflammatory pathway, triggered by acetylcholine released from the efferent vagus nerve endings [26, 27], and are sensitive to endogenous acetylcholine produced by immune cells [28]. In addition, the α7β2 nAChRs located in mitochondria regulate the mitochondrial apoptosis pathway and, therefore, influence the viability of brain cells [29].

Importantly, α7 nAChRs can bind soluble Aβ with very high affinity [30]. The formed Aβ-α7 nAChR complexes are then endocytosed, leading to fewer membrane-bound α7 nAChRs and the intracellular accumulation of amyloid [31]. Indeed, several studies have demonstrated a profound loss of nicotinic acetylcholine receptors in the postmortem patient’s brains (reviewed in ref. [31]). The data specifically focusing on α7 nAChRs is, however, less consistent. Thus, expression of α7 nAChRs was shown to increase in hippocampal and cortical astrocytes from AD patients but to decrease in hippocampal and cortical neurons [32, 33]; and both deletion [34, 35] and stimulation [36] of α7 nAChRs were shown to improve cognition in different mouse models of AD. Moreover, several (partial) agonists of α7 nAChRs were tested without big success in clinical trials [31], which, however, were mostly of short duration. This calls for a better understanding of the underlying molecular mechanisms.

To understand when the α7 nAChR dysfunction occurs during AD development, which molecular mechanisms are involved and how one can alleviate this dysfunction, we analyzed APPswe/PS1_G384A_ mice, overexpressing human amyloid precursor protein (APP) with the Swedish double mutation (K670N, M671L) and a mutant presenilin 1 (PS 1, G384A mutation) under the control of Thy-1 promoter [10].

## 2. Materials and methods

### 2.1. Materials

All reagents were of chemical grade and were purchased from Sigma-Aldrich (Saint Louis, USA) unless otherwise indicated. Antibodies against α7(1-208) [37], α3(181-192), α4(181-192), α7(179-190) [38], α9(11-23) [39], β2(190-200) or β4(190-200) [40] nAChR fragments and cytochrome *c* [41] were previously produced, characterized and biotinylated in the Kyiv laboratory. Mouse IL-1β antibody pair (ab210895), IL-6 antibody pair (ab213749) and IL-10 antibody pair (ab214473) were from ABCAM. Antibodies against Aβ_1-42_ (cat # 44-3449) were from Invitrogen.

### 2.2. Animals

All experimental procedures were performed following institutional animal welfare guidelines and approved by the state government of Baden-Württemberg, Germany. 1.5 and 6 months old C57BL/6N (wild type, WT) or APPswe/PS1G_384A_ mice (AD) mice of either sex were used in this study. Animals were housed under standard conditions with a 12-hour light/dark cycle and free access to food and water. Females stayed in groups of 3-5 mice, males were kept individually. All procedures complied with the [ARRIVE guidelines](https://www.nc3rs.org.uk/arrive-guidelines) and were carried out following the [EU Directive 2010/63/EU for animal experiments](https://environment.ec.europa.eu/topics/chemicals/animals-science_en).

### 2.3. Behavioral experiments

In the Novel Object Recognition (NOR) test [42] after a 10-min-long habituation mice were allowed to explore two identical objects for 10 min and then, after a 15-20 min break, one object was replaced by a novel object of similar size but different shape and color and the number of explorations of the two objects as well as the total exploration time spent with the given object were measured during the subsequent 10-min-long session (Fig. 1A). The results of the NOR test are presented as Discrimination Index (DI) I and II. DI I was calculated as the difference in the total exploration time of “novel” (T1) or “familiar” (T2) objects divided by the total time of exploration of both objects [DI I= (T1 − T2)/(T1 + T2)]. DI II was calculated as the difference in the number of contacts with “novel” (N1) or “familiar” (N2) objects divided by the total number of contacts with both objects [DI II= (N1 − N2)/(N1 + N2)]. The lack of novel object preference, expressed as a DI decrease, was interpreted as episodic memory impairment.

**Fig. 1.**
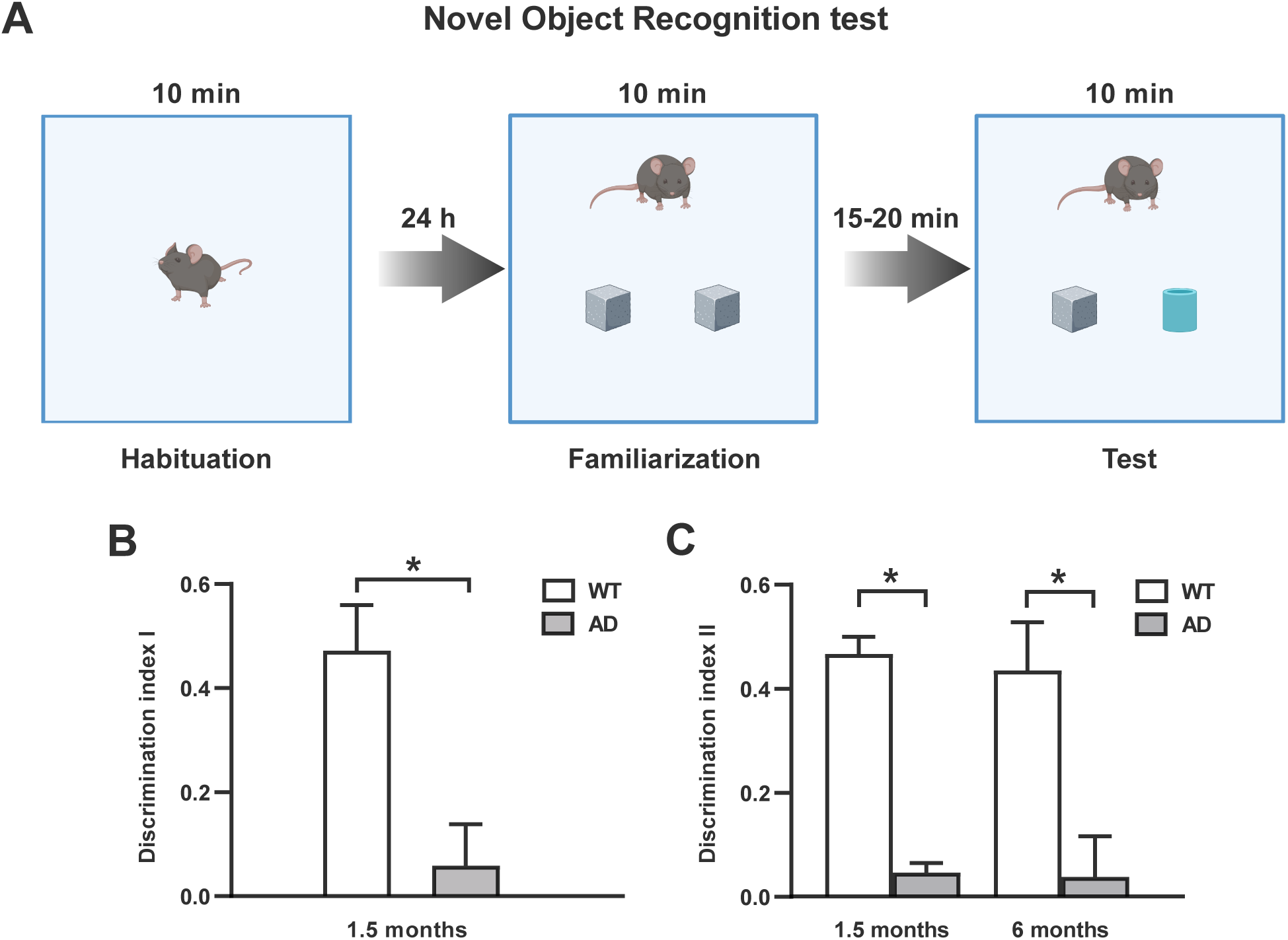
Impairment of the episodic memory of the 1.5 and 6 months old WT and AD mice. (**A**) Schematic of experimental arrangement in the NOR test (created with BioRender.com, see Materials and methods for details). (**B**, **C**) Bar graphs, illustrating the Discrimination indexes I (n=8 WT and 15 AD mice, Student’s t-test, p=0.004) and II (WT 1.5 months old, n=9; 6 months old, n=4 mice; AD: 1.5 months old, n=12; 6 months old, n=5 mice; Two way ANOVA followed by Bonferroni’s multiple comparisons test, *p<0.05 for all comparisons), calculated as described in Materials and methods.

In the open field test, enabling to characterize explorative and locomotor activity [43], the animals were individually placed in a rectangular (45 × 45 cm) novel open field and observed for 10 min. The locomotor activity was evaluated as the total distance traveled and the average speed. At the end of the experiment, the animals were decapitated under deep CO_2_ anesthesia, the brains were removed, homogenized in PBS or the mitochondria isolation buffer (containing 10 mM HEPES, 1 mM EGTA, 200 mM saccharose, pH 7.4, t=4°C) and processed as described below.

### 2.4. Cytochrome c release from mitochondria

Mitochondria were isolated from mouse brain homogenates by differential centrifugation according to standard published procedures [44]. The freshly isolated mitochondria (120 µg of protein per ml), resuspended in 10 mM HEPES, 125 mM KCl, 25 mM NaCl, 5 mM sodium succinate and 0.1 mM Pi(K), pH 7.4, were incubated with either 0.09 µM CaCl_2_ or 0.5 mM H_2_O_2_ for 5 min at room temperature and were immediately pelleted by centrifugation (10 min, 7000 g) at 4°C. The supernatants were tested for the presence of cytochrome *c* (Cyt *c*) by sandwich ELISA assay, while the pellets were frozen for further investigation for the presence of Cyt c, nAChR subunits and Aβ_1-42_ as described below.

### 2.5. ELISA assays

The pellets of homogenized brains or mitochondria were frozen at -20 °C in the lysing buffer (0.01 M Tris-HCl, pH 8.0; 0.14 NaCl; 0.025% NaN_3_; 1% Tween-20 and protease inhibitors cocktail). After being thawed, they were lysed for 2 h on ice upon intensive stirring. The resulting lysates were pelleted by centrifugation (20 min at 20000 g). The protein concentration was established with the BCA Protein Assay kit (#23227, Thermo Fisher Scientific, Rockford, USA).

The levels of nAChR subunits and Aβ_1-42_ bound to α7 nAChR in the brain or mitochondria preparations were determined as described in [45]. Briefly, the detergent lysates of either the whole brain or mitochondria were applied into the wells of Nunc Maxisorp immunoplates (1 µg of protein per 0.05 ml per well) coated with rabbit α7(1-208)-specific antibody (20 µg/ml). The bound subunits were detected with the second biotinylated α3(181-192)-, α4(181-192)-, α7(179-190)-, α9(11-23), β2(190-200)- or β4(190-200)-specific antibody, while the bound Aβ_1-42_ was detected with biotinylated Aβ_1-42_-specific antibody.

The level of free Aβ_1-42_ was revealed using polyclonal Aβ_1-42_-specific antibody (#44-3449, Thermo Fisher Scientific), a portion of it being biotinylated. The plates were coated with non-biotinylated antibody (20 µg/ml), while Aβ_1-42_ bound from the brain or mitochondria detergent lysate was detected with biotinylated antibody. The bound biotinylated antibodies were visualized with Neutravidin-peroxidase conjugate and *o*-phenylenediamine-containing substrate solution. To measure the levels of α7-bound Aβ1-42, the plates were coated with the antibodies against the whole extracellular domain of the α7 subunit α7(1-208), while the moiety bound from the brain or mitochondria detergent lysate was revealed with Aβ1-42-specific antibody. The resulting OD value was normalized to that of the α7-specific value obtained in the sandwich of α7(1-208) and α7(179-190)-specific antibodies.

To detect Cyt *c* released from mitochondria, the plates were coated with Cyt *c*-specific rabbit polyclonal antibody [41] and blocked with 1% BSA. Either the mitochondria supernatants (non-diluted) or detergent lysates of mitochondria pellets (1 µg of protein per 0.05 ml per well) were applied to the wells with adsorbed antibody for 2 h at 37°C and, after extensive washing, the biotinylated anti-Cyt *c* antibody was applied for additional 1h to be further revealed with Neutravidin-peroxidase conjugate and o-phenylenediamine-containing substrate solution.

In Aβ_1-42_- and Cyt *c*-specific assays, we used similar antibodies for coating and detection, because, due to their polyclonal nature, they recognized multiple epitopes on the Aβ_1-42_ or Cyt *c* molecule; such an approach was shown to be efficient in many previous experiments [41, 45, 46]. The cytokine levels (IL-1β, IL-6 and IL-10) were measured by Sandwich ELISA according to recommendations of the kit’s manufacturer (Abcam, UK): Mouse IL-6 Matched Antibody Pair Kit (# ab213749), Mouse IL-1 beta Matched Antibody Pair Kit (#ab210895) and Mouse IL-10 ELISA Set (#ab47599). The optical density was read at 490 nm or 450 nm (for the cytokines) using Stat-Fax 2000 ELISA Reader (Awareness Technologies, USA). All ELISA assays were performed in triplicates and mean values were used for statistical analyses.

### 2.6. Statistical analysis

Statistical analyses were performed using the GraphPad Prism (Version 9.5.1). The normality of data distribution was tested with the Shapiro–Wilk test. For normally distributed data, the Student’s t-test or ANOVA followed by Tukey’s or Bonferroni’s multiple comparison test was used. The two-way ANOVA followed by either the Turkey’s or Bonferroni’s test for multiple comparisons, was used to compare more than two groups and factors. *p values < 0.05 were considered statistically significant. Unless otherwise indicated, all data are shown as mean ± SEM.

## 3. Results

### 3.1. Early impairment of episodic memory, α7 nAChR content and mitochondrial stability in APPswe/PS1G384A mice

In the first set of experiments, we compared WT and APPswe/PS1_G384A_ (AD) mice of two different ages: 1.5 months (the stage before the deposition of dense core amyloid plaques [10]), and 6 months (stage of widespread amyloid deposition and inferior performance in discriminatory water maze and Y-mase tests [10]). Surprisingly, as documented by the NOR test, the AD mice demonstrated worse episodic memory compared to WT already at 1.5 months and this deficit was maintained at 6 months of age (Fig. 1).

Subsequently, the brain preparations of all groups of mice were examined for the levels of cytokines, nAChR subunits and Aβ_1-42_. The WT data revealed similarly low levels of IL-1β in 1.5 and 6 months old mice; a small but significant aging-associated decrease in the level of IL-6 and a small but significant increase in the level of IL-10 (Fig. 2). The brains of 1.5 months old AD mice contained significantly more pro-inflammatory cytokines IL-1β and IL-6 and similar levels of an anti-inflammatory cytokine IL-10 compared to age-matched WT mice. In 6 months old AD mice, the levels of IL-1β decreased significantly compared to 1.5 months old AD mice and were similar to that measured in WT littermates of the same age, while the levels of IL-6 and IL-10 were higher in AD compared to WT mice (Fig. 2). Overall, the levels of pro-inflammatory cytokines (IL-1β and IL-6) decreased, and the levels of an anti-inflammatory IL-10 increased with disease progression, thus revealing an early pro-inflammatory brain state in 1.5 months old AD mice, which was partially attenuated at 6 months of age.

**Fig. 2.**
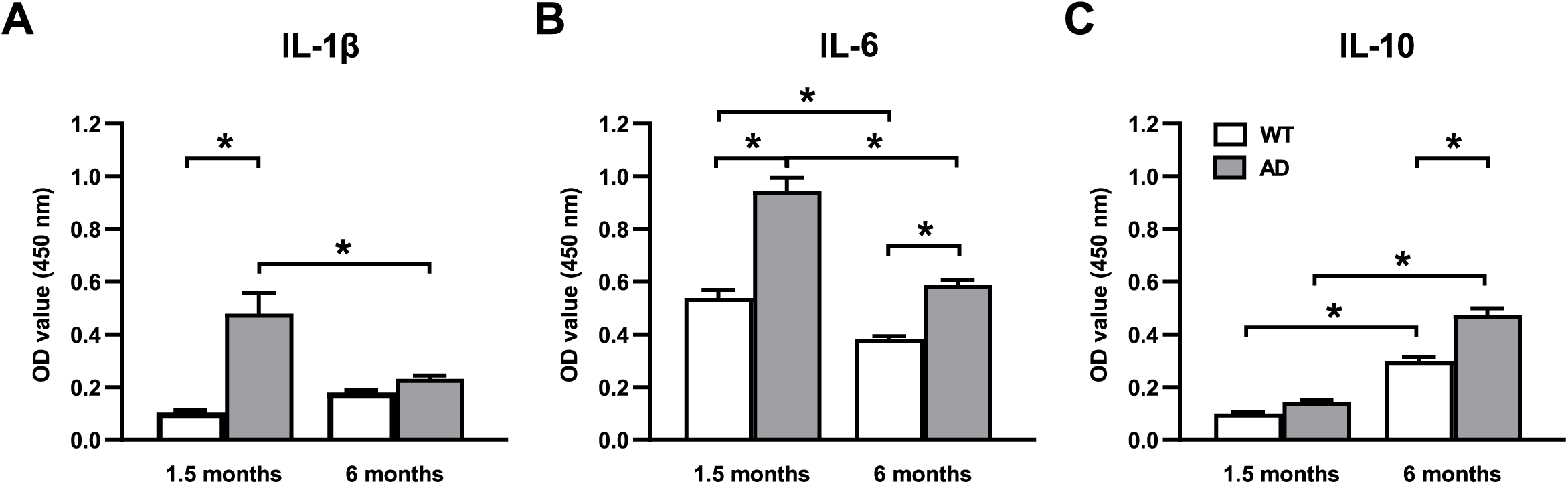
Levels of cytokines in the brain detergent lysates of the 1.5 and 6 months old WT or AD mice. (**A**-**C**) Bar graphs, illustrating the cytokine levels (IL-1β, IL-6 and IL-10), measured by Sandwich ELISA according to the manufacturer’s protocol (n=4 mice per group; two-way ANOVA followed by Tukey’s multiple comparisons test; *p<0.05 for all comparisons).

The brains of AD mice contained significantly fewer α7 nAChR subunits compared to the brains of WT mice at both 1.5 and 6 months of age (Fig. 3A, B). The levels of other subunits were also non-identical: α3 and α4 subunits were downregulated whereas β4 subunits were upregulated at 1.5 months of age. At 6 months of age, β4 subunits were downregulated and α3, as well as α9 subunits, were upregulated in AD compared to WT mice. Similarly, at 1.5 months of age, the brain mitochondria of AD mice contained significantly fewer α7 and β2 nAChRs and more β4 nAChRs compared to the mitochondria of WT mice. The α7 deficiency became more pronounced in the mitochondria of 6 months old mice, in which α9 and β2 nAChRs were also downregulated (Fig. 3C, D). Of note, there was very little change in the level of nAChRs between 1.5 and 6 months old WT mice for both brain and brain mitochondria (Figure S1A and B), whereas the age-related changes in the AD group (Figure S1C and D) reflected a decrease in the pro-inflammatory brain state, consistent with data shown in Fig. 2. Together, these data reveal a persistent down-regulation of α7 nAChRs in APPswe/PS1G_384A_ mice starting from 1.5 months of age, while the levels of other nAChR subunits vary with age and disease progression.

**Fig. 3.**
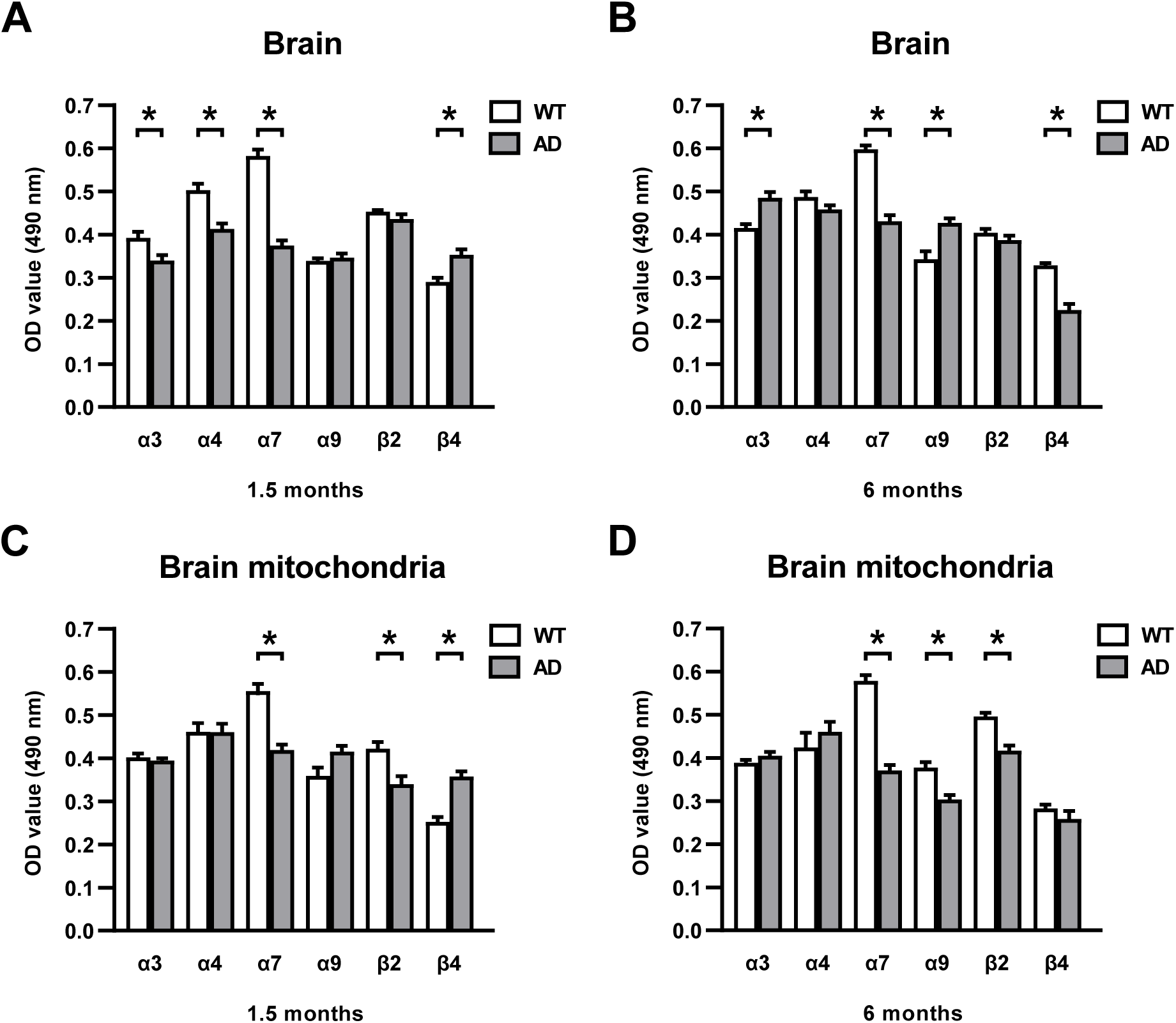
The levels of nAChR subunits in the brain or brain mitochondria of the WT and AD mice. (**A**, **B**) Bar graphs, illustrating the levels of different nAChR subunits in the detergent lysates of the whole brain of 1.5 (**A**) and 6 (**B**) months old WT and AD mice. (**C**, **D**) similar analyses as in (**A**, **B**) but conducted in the detergent lysates of brain mitochondria (n=4 mice per group; two-way ANOVA followed by Bonferroni’s multiple comparisons test; *p<0.05 for all comparisons).

Next, we measured the levels of α7-bound Aβ_1-42_ in either the brain or brain mitochondria of AD compared to WT mice. As shown in Fig. 4, both the whole brain and the mitochondria preparations of AD mice contained significantly more Aβ_1-42_ coupled to α7 than those of WT mice. For brain mitochondria, this ratio further increased from 1.5 to 6 months of age.

**Fig. 4.**
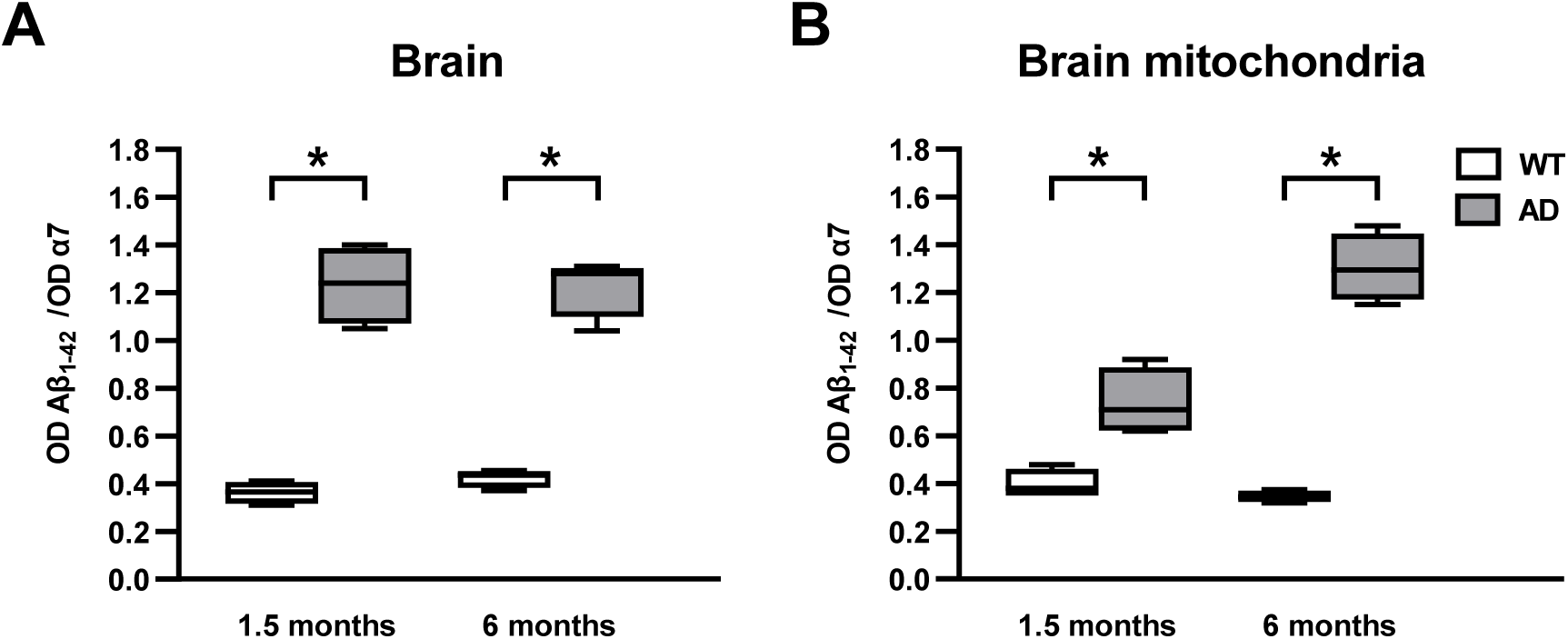
The levels of α7-bound Aβ_1-42_ in the brain and brain mitochondria of the WT and AD mice. (**A**, **B**) Box plots, illustrating the levels of α7-bound Aβ_1-42_ in the detergent lysates of either the whole brain (**A**) or brain mitochondria (**B**) (n=4 mice per group; two-way ANOVA followed by Bonferroni’s multiple comparisons test; *p<0.05 for all comparisons). Data are shown as median ± IQR. Here and below boxes extend from the 25th to 75th percentiles; the whiskers cover the whole data range.

Finally, the freshly isolated brain mitochondria of either WT or AD mice were compared in a cytochrome *c* (Cyt *c*) release assay in control (untreated mitochondria) and under treatment with either 0.9 µM Ca^2+^ or 0.5 mM H_2_O_2_. As shown in Fig. 5, the mitochondria of 6 months old AD mice released significantly higher amounts of Cyt *c* compared to mitochondria of WT mice even without Ca^2+^ or H_2_O_2_ treatment (control), demonstrating their decreased stability compared to WT mitochondria. The difference became more pronounced under Ca^2+^ or H_2_O_2_ treatment and was seen both in 1.5 and 6 months old mice. Because Cyt *c* release from mitochondria is a signal initiating the mitochondrial apoptosis pathway [47], these data suggest that the mitochondria of AD mice are more susceptible to reactive oxygen species- or Ca^2+^-induced apoptogenic stimuli. This is especially important given the well-known AD-mediated enhancement of ROS production and ongoing Ca^2+^ signaling throughout the brain [6, 48, 49].

**Fig. 5.**
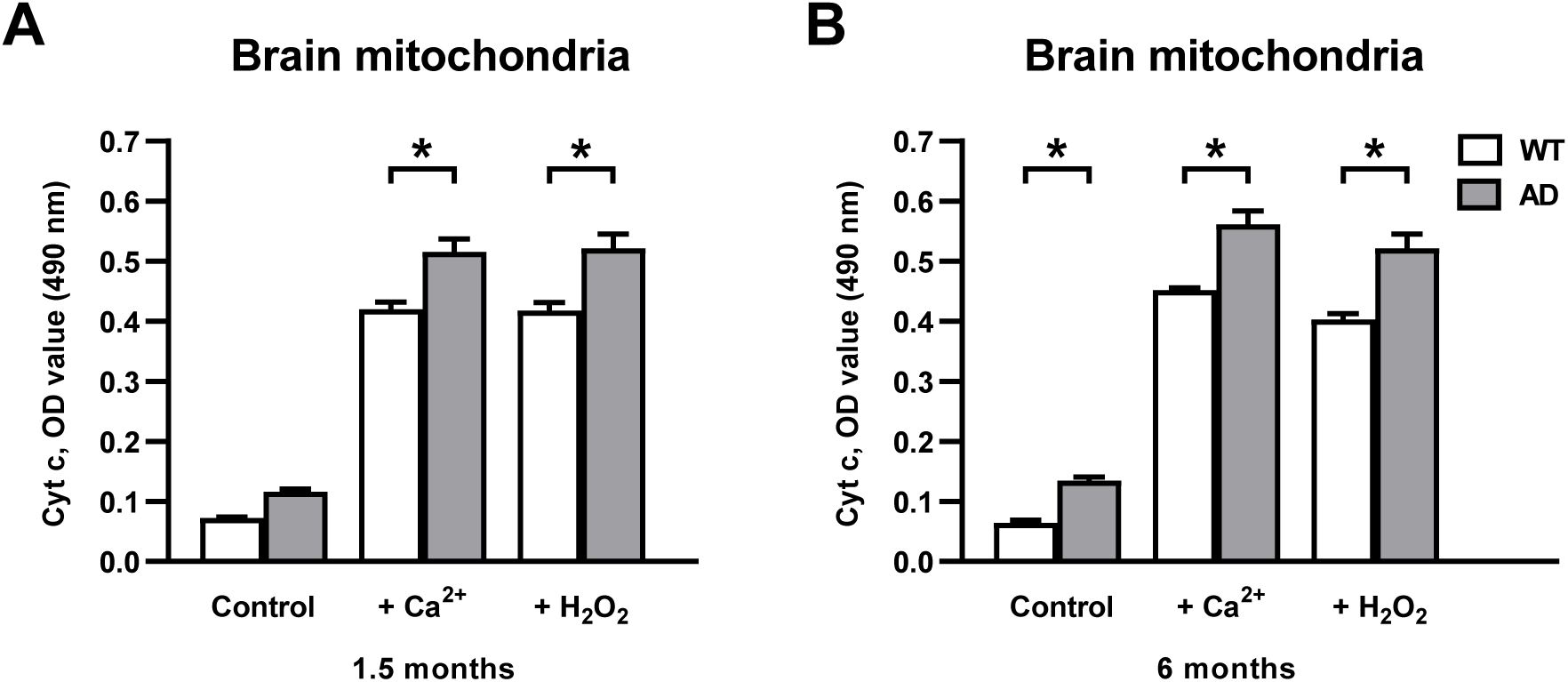
Cytochrome *c* released from the brain mitochondria of WT and AD mice in control or in response to either Ca^2+^ or H_2_O_2_. (**A**, **B**) Bar graphs, illustrating the levels of Cyt *c*, measured by sandwich ELISA assay in the respective supernatants (see Materials and methods for details) of either 1.5 (**A**) or 6 (**B**) months old WT and AD mice (n=4 mice per group; two-way ANOVA followed by Bonferroni’s multiple comparisons test; *p<0.05 for all comparisons).

Taken together, the above data show that APPswe/PS1G_384A_ mice differ from the age-matched WT controls already at 1.5 months of age. Their brains contain increased levels of pro-inflammatory cytokines, decreased levels of α7 nAChRs and increased amounts of α7-bound Aβ_1-42_, along with an impaired cortex-dependent episodic memory and increased risk of apoptic cell death.

### 3.2. The effects of α7 nAChR stimulation in 1.5 months old APPswe/PS1G_384A_ mice

To test whether improving the α7 nAChR-mediated signaling can alleviate the described above amnestic state, we treated 1.5 months old APPswe/PS1G_384A_ mice with a selective α7 nAChR agonist PNU282987 [50]. The mice received daily i.p. injections of either PNU282987 in PBS (5 mg/kg, as in [51]) or PBS alone for 7 consecutive days and were tested in NOR *and* open field behavioral tests immediately before the treatment and at the end of treatment (on day 8). The treatment was well tolerated, as the animals (n=6 in each group) did not lose weight (median±IQR: 15.5±6.0/17.0±5.0 g before/after the treatment with PBS and 17.7±5.3/ 18.0±4.4 g with PNU282987 (Wilcoxon matched-pair signed rank test, p=0.31 and 0.06, respectively)) or showed any behavioral abnormalities. Two different readouts (the number of explorations and the total exploration time, see Materials and methods) were used to judge the animal’s performance in the NOR test. In the total exploration time, the PBS-treated mice showed a small downward trend toward memory impairment (Fig. 6A). This trend was successfully prevented by the treatment with PNU282987 in all but one animal (Fig. 6B, C). The memory-enhancing effect of PNU282987 was more visible when analyzing the number of explorations (Fig. 6D-F), with PNU282987-treated AD mice showing improved episodic memory compared to mice receiving PBS injections. The PNU282987-induced improvement in cognitive performance was not directly related to changes in motor function, as there were no alterations in locomotor or explorative activity between WT and AD mice in response to PNU282987 treatment (total distance traveled: 2.67±0.36 for PBS and 2.81±0.36 for PNU282987 treatment, Student’s t-test p=0.78; average speed: 4.45±0.59 for PBS and 4.69±0.60 for PNU282987 treatment, Student’s t-test p=0.78; n=9 and 8 mice, respectively).

**Fig. 6.**
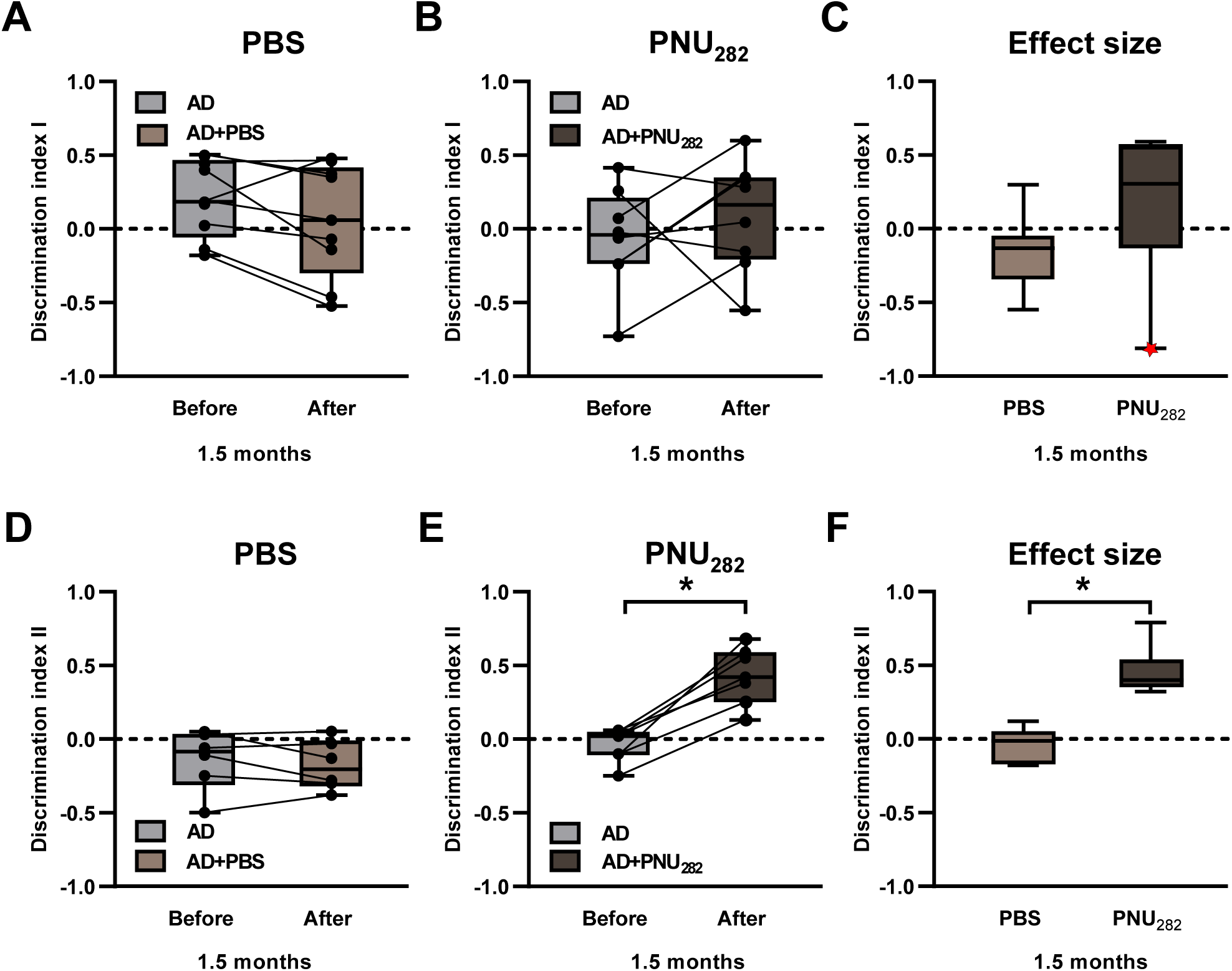
Assessment of the episodic memory of the 1.5-month-old AD mice before and after a week-long treatment with either PBS or PNU282987. (**A**, **B**) Box plots showing Discrimination indexes I (n=9 and 8 mice; p=0.08 and 0.41, respectively, Paired t-test) measured in the NOR test in mice, treated with either PBS (**A**) or PNU282987 (**B**). (**C**) Box plots, illustrating the effect size, calculated by subtracting, for each mouse, the Discrimination index measured after the treatment from the Discrimination index, measured before the treatment. The red dot marks the data point, most distant to the median but not an outlier, which prevents the difference from becoming significant (Student’s t-test, p=0.11). (**D**-**F**) the same analyses as in (**A**-**C**), conducted for the Discrimination index II. **D**: n=6 mice, Paired t-test, p=0.47; **E**: n=7 mice, Paired t-test, p=3.0x10^-4^; F: Student’s t-test, p <10^-4^.

We again confirmed that the brains of APPswe/PS1G_384A_ mice contained significantly lower levels of α7 nAChRs compared to WT mice. However, their level was increased by the PNU282987 treatment (Fig. 7A). The β2 subunits were also downregulated in AD and upregulated back to the control level in the PNU282987-treated animals. Similarly, PNU282987 treatment counteracted the AD-induced changes in the level of β4 subunits. The levels of other subunits, which were different between WT and APPswe/PS1G_384A_ mice, were not affected by PNU282987. In this experiment, we also measured the levels of Aβ_1-42_ either free or coupled to α7. The brains of APPswe/PS1G_384A_ mice contained significantly more free Aβ_1-42_ compared to the brains of WT mice (Fig. 7C), as well as higher levels of α7-coupled Aβ_1-42_ (Fig. 7E). Whereas the level of free Aβ_1-42_ was not affected by PNU282987 (Fig. 7C), PNU282987 treatment significantly decreased the level of α7-coupled Aβ_1-42_ (Fig. 7E).

**Fig. 7.**
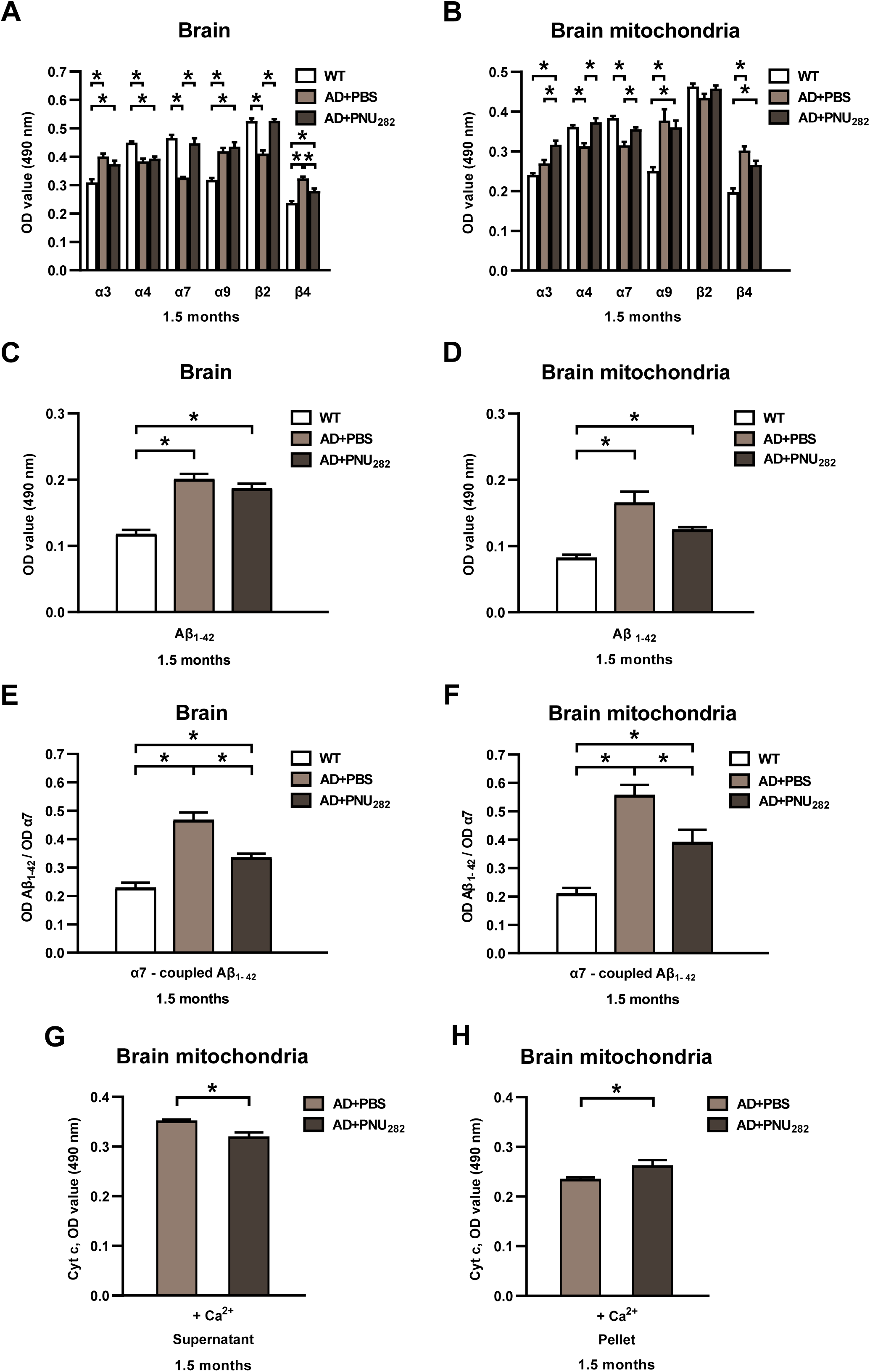
Effect of the week-long PNU282987 treatment on the expression of nAChR subunits, Aβ levels in, and cytochrome *c* release from the brain and brain mitochondria. (**A, B**) Bar graphs illustrating the levels of different nAChR subunits, measured simultaneously in the whole brain (**A**) or brain mitochondria (**B**) detergent lysates of 1.5-month-old WT and AD mice, treated either with PBS or PNU282987, as indicated in the insert. Two-way ANOVA followed by Bonferroni’s multiple comparisons test, *p<0.05 for all comparisons (n=6, 6, 7 WT, AD+PBS and AD+PNU282 mice, respectively). (**C, D**) Bar graphs illustrating the levels of free soluble Aβ_1-42_ in the detergent lysates of the whole brain (**C**) or brain mitochondria (**D**) of 1.5-month-old WT mice or AD mice, treated either with PBS or PNU282987 (**C**: n=5, 5, 7; **D**: n=6, 6, 7 WT, AD+PBS and AD+PNU282 mice, respectively). One-way ANOVA without (**C**) or with Welch’s correction for unequal variances (**D**) followed by Bonferroni’s or Dunnett’s T3 multiple comparisons tests, respectively, *p<0.05 for all comparisons. (**E**, **F**) Bar graphs, illustrating the levels of α7-bound Aβ_1-42_ in the detergent lysates of either the whole brain (**E**) or brain mitochondria (**F**) (n=6 mice per group). One-way ANOVA followed by Bonferroni’s multiple comparisons test, *p<0.05 for all comparisons. (**G**, **H**) Bar graphs illustrating the levels of Ca^2+^-induced Cyt *c* either released into the supernatant (**G**) or remaining in the mitochondria pellet (**H**) (n=6, 7 AD+PBS and AD+PNU282 mice, respectively; Unpaired t-test with Welch’s correction p=8.4x10^-3^ (**G**) and 4.7x10^-2^ (**H**)).

Consistent with the data obtained in the whole brain preparation, PNU282987 increased the levels of α7 subunits in the mitochondria of APPswe/PS1G_384A_ mice (Fig. 7B). Besides, the levels of α3 and α4 subunits increased significantly after the PNU282987 treatment while the levels of other subunits were not affected. This may mean that, in addition to α7 nAChRs, PNU282987 favored the up-regulation of the α3/α4-containing nAChR subtype. The levels of free (Fig. 7D) and α7-coupled (Fig. 7F) Aβ_1-42_ were higher in mitochondria of APPswe/PS1G_384A_ compared to WT mice and were slightly (Fig. 7D) or significantly (Fig. 7F) decreased by PNU282987. Furthermore, the level of Cyt *c*, released from isolated mitochondria of APPswe/PS1G_384A_ mice into the supernatant in response to Ca^2+^, was reduced by PNU282987 (Fig. 7G). Correspondingly, the content of Cyt *c* within the mitochondria pellet increased upon PNU282987 treatment (Fig. 7H).

### 3.3. Disease-modifying effect of prolonged stimulation of α7 nAChRs

To test the therapeutic potential of α7 nAChRs stimulation, we treated the APPswe/PS1G_384A_ mice once a week with i.p. injections of either PNU282987 in PBS (5 mg/kg) or PBS alone for 17 consecutive weeks (chronic treatment regimen). Over the treatment time, we monitored the behavior and well-being of mice in the control and experimental groups and did not observe any gross differences. Both groups continuously gained weight and showed similar behavior in the open field test at the end of treatment (Figure S2). When tested in the NOR test immediately before and at the end of chronic treatment, only the PNU282987-treated mice improved significantly (Fig. 8A-F), reaching the level of performance, comparable to that of WT mice (n=4 WT and 6 PNU282987-treated AD mice, Student’s t-test, p=0.08). In addition, the chronic treatment regimen decreased the amount of IL-6 and significantly increased the amount of IL-10 in the brain, compared to that of PBS-treated mice (Fig. 8G, H).

**Fig. 8.**
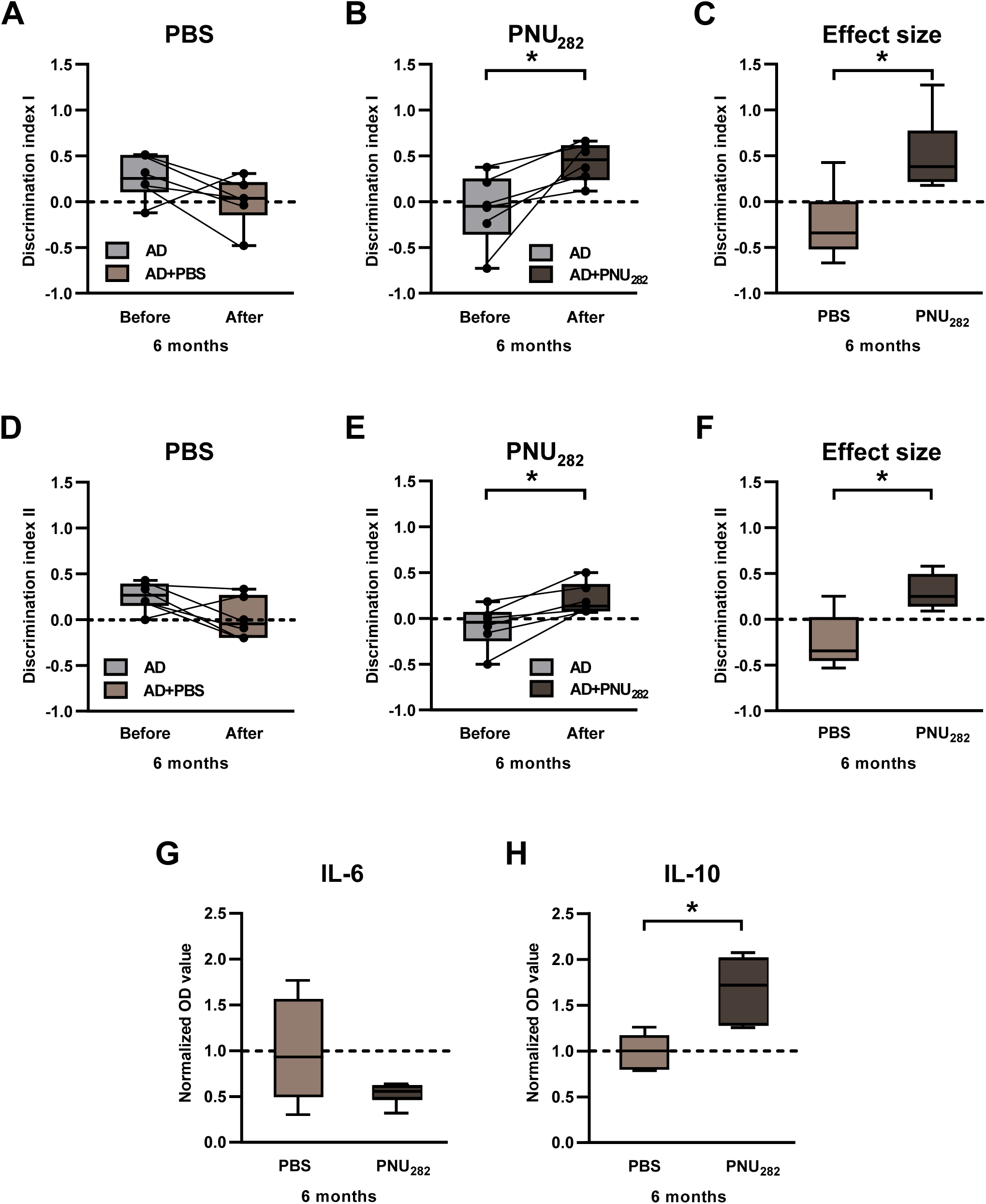
Episodic memory and the levels of cytokines in the brain detergent lysates of AD mice before and after a 4-month-long PNU282987 treatment. (**A**, **B**) Box plots showing Discrimination indexes I (n=6 mice per group; p=0.16 and 2.8x10^-2^, respectively, Paired t-test) measured in the NOR test in mice, treated with either PBS (**A**) or PNU282987 (**B**). (**C**) Box plots, illustrating the effect size, calculated by subtracting, for each mouse, the Discrimination index measured after the treatment from the Discrimination index, measured before the treatment (Student’s t-test, p=7.1x10^-3^). (**D**-**F**) the same analyses as in (**A**-**C**), conducted for the Discrimination index II. **D**: n=6 mice, Paired t-test, p=0.1; **E**: n=6 mice, Paired t-test, p=0.01; **F**: Student’s t-test, p=3.4x10^-3^. (**G**, **H**) Box plots, illustrating the levels of IL-6 (**G**) and IL-10 (**H**), measured by Sandwich ELISA and normalized to the mean value measured under the PBS treatment (n=6 mice per group; Wilcoxon rank sum test, p=0.31 and 4.3x10^-3^, respectively).

Furthermore, the chronic treatment significantly increased the levels of all nAChRs in the brain and all but α3 subunit in brain mitochondria (Fig. 9A-B). Next to decreasing the levels of α7-bound Aβ_1-42_ in the brain and brain mitochondria, the chronic treatment also significantly decreased the brain levels of free Aβ_1-42_ (Fig. 9C-F). Moreover, the level of Cyt *c*, released from isolated mitochondria into the supernatant in response to Ca^2+^ was reduced, and the content of Cyt *c* within the mitochondria pellet was increased upon the chronic PNU282987 treatment (Fig. 9G, H), with both effects being much stronger compared to a week-long-treatment (Fig. 7G, H).

**Fig. 9.**
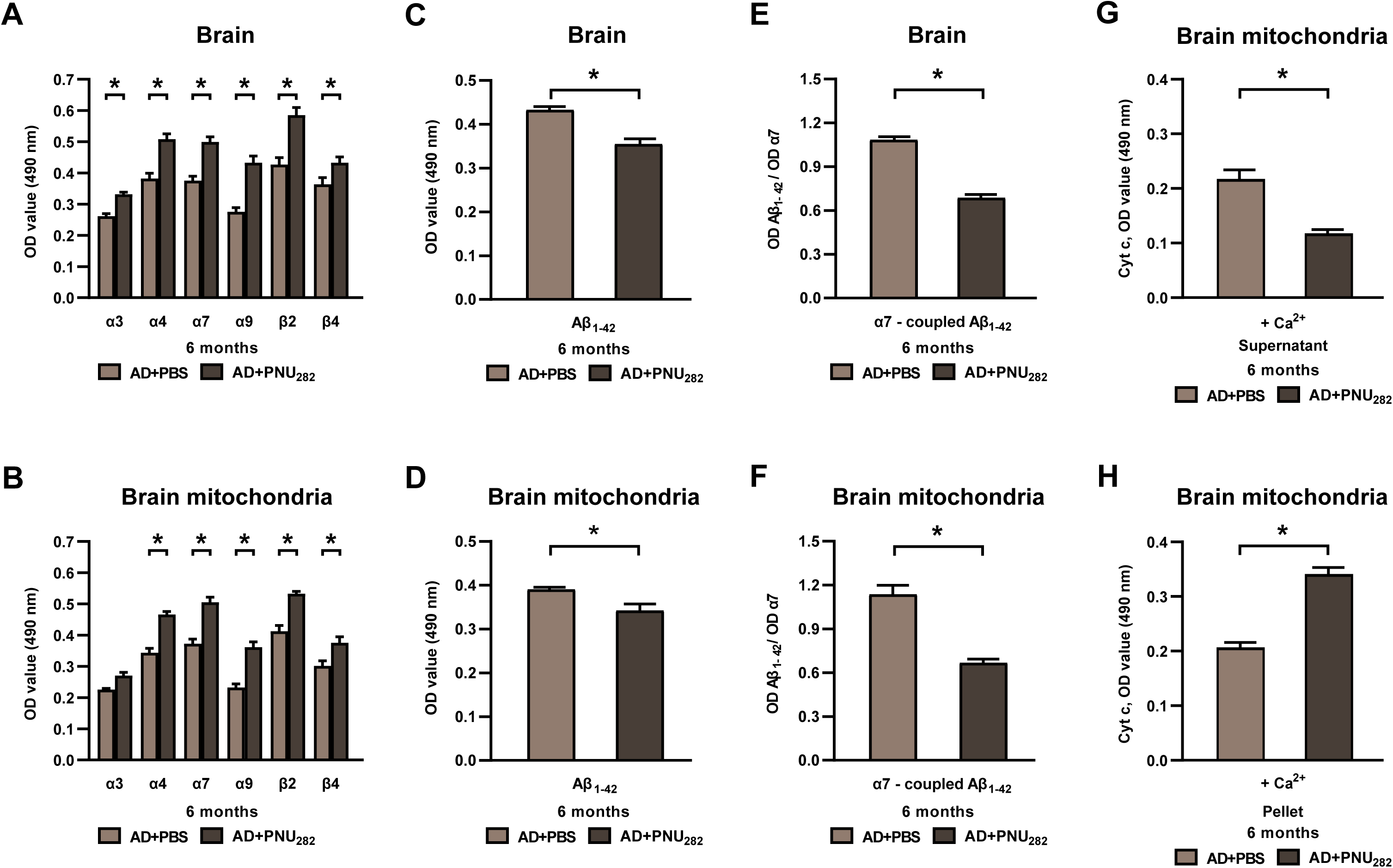
Effect of the 4-month-long PNU282987 treatment on the expression of α7 nAChRs, Aβ levels and cytochrome *c* release from brain mitochondria. (**A**, **B**) Bar graphs, illustrating the levels of different nAChR subunits in the detergent lysates of the whole brain (**A**) or brain mitochondria (**B**) of AD mice, treated over 4 months with either PBS or PNU282987. (**C**-**F**) Bar graphs illustrating either free soluble Aβ_1-42_ levels in the brain (**C**) and brain mitochondria (**D**) or levels of α7-coupled Aβ_1-42_ in the brain (**E**) or brain mitochondria (**F**) of AD mice treated with either PBS or PNU282987. **A**-**C** and **E**-**F**: Student’s t-test, *p <0.001 for all comparisons; **D**: Student’s t-test with Welch’s correction for unequal variances, p=2.3x10^-2^, n=6 mice per group. (**G**, **H**) Bar graphs illustrating the levels of Ca^2+^-induced Cyt *c* either released into the supernatant (**G**) or remaining in the mitochondria pellet (**H**). Student’s t-test p=3x10^-^4 (**G**) and p=8.8x10^-6^ (**H**), n=6 mice per group.

Thus, the chronic PNU282987 treatment during the disease development phase (1.5-6 months of age) successfully reduced the intra- (α7-bound Aβ_1-42_) and extracellular (free Aβ_1-42_) accumulation of amyloid, as well as disease-mediated (i) downregulation of nAChRs, (ii) Ca^2+^-induced release of apoptogenic stimuli from mitochondria, (iii) neuroinflammation and (iv) memory impairment, thus underlining the power of α7 nAChR modulation for disease-modifying treatment.

## 4. Discussion

Multiple evidence suggests the involvement of brain α7 nAChRs in the development and progression of AD (reviewed in refs. [31, 52, 53]). However, the time course of α7 dysfunction during AD development and the underlying molecular mechanisms are not completely understood. By using a well-characterized mouse model of AD, here we show that α7 nAChRs are downregulated at the very early stages of AD pathology and their activation with an agonist can improve multiple disease-related dysfunctions in the brain and brain mitochondria of young mice as well as counteract the development of these symptoms with age.

The data obtained revealed a strong and unexpectedly early (at 1.5 months of age) impairment of episodic memory, which is mainly perirhinal cortex-dependent [54], along with the pro-inflammatory brain state (2-4-fold increase in levels of IL-1β and IL-6) and strong and consistent downregulation of α7 nAChRs within the brain. This is unexpected, since in APPswe/PS1_G384A_ mice at this age (pre-depositing disease phase) there is no plaque accumulation throughout the cortex (and the entire brain), the amount of soluble Aβ_1-40_ and Aβ_1-42_ is very low, and the cortical neural network activity is still normal [10]. The only abnormality, known so far for pre-depositing APPswe/PS1_G384A_ mice, is the profound hyperactivity of the hippocampal pyramidal neurons, likely triggered by soluble Aβ oligomers [55]. Despite this hyperactivity, however, the animals’ performance in the hippocampus-dependent discriminatory water maze task as well as in the hippocampus, septum, basal forebrain, and prefrontal cortex-dependent Y-maze working memory test remains normal [10].

We did observe, however, enhanced binding of the soluble Aβ oligomers to α7 nAChR in the pre-depositing phase both in the brain and brain mitochondria (Fig. 4). Moreover, within the brain the amount of Aβ/α7 nAChR complexes saturated already at 1.5 months of age and did not increase further during disease progression. According to the literature, Aβ binds within an orthosteric binding site of α7 nAChR and can be displaced by α7-selective agonists, competitive antagonists, or even positive allosteric modulators [56]. Moreover, in the presence of Aβ α7 undergoes concentration-dependent conformational changes that bring it toward the desensitized state. It was shown that Aβ directly affects α7 function by acting as an agonist and a negative modulator. It activates α7 nAChR at low concentrations, while in the presence of higher Aβ concentrations or prolonged exposure, it reduces α7 nAChR activity. This might contribute to deficits in cholinergic signaling as well as the internalization of Aβ/α7 nAChR complexes [31, 56, 57]. The internalized Aβ oligomers, in turn, were recently shown to directly induce neuronal hyperactivity [58], a known hallmark of AD [6, 10]. In fact, intracellular accumulations of Aβ are indeed visible in neurons from deeper cortical layers in the 1.6-month-old APPswe/PS1_G384A_ mice (see Fig. S1F in [10]). Furthermore, our data show that even in pre-depositing AD mice the brain mitochondria release more Cyt *c* under control conditions and are more susceptible to the effect of apoptogenic agents, thus paving the way to neurodegeneration.

So far, only amyloid-depositing mice were treated with the α7 nAChR agonists (A-582941 or PNU282987) in preclinical studies [36, 59, 60]. A week-long daily i.p. treatment of 6- or 10-month-old APP/PS1 mice, for example, significantly improved animals’ performance in the Morris water maze test, along with the significant upregulation of synaptic proteins (e.g., SYN, PSD95, SNAP25, DYN1, AP180), an increase in the number of hippocampal synapses, downregulation of the Aβ production machinery (BACE1, BACE2, ADAM10) and a decrease in the level of oxidative stress. The effect on amyloid deposition was less conclusive, likely pointing to a decrease in the intracellular Aβ without a reduction in the density of dense core plaques [59, 60]. In another study, a 3-month-long oral treatment of aged (15-month-old) 3xTg-AD mice with robust plaques and tangles significantly improved animals’ performance in the Morris water maze, novel object recognition and contextual fear memory tests, without any changes in the Aβ production (APP, C99/C83, ADAM10, ADAM17, BACE1) or degradation (insulin-degrading enzyme, neprilysin) machinery, the extent of neuroinflammation, tau phosphorylation or levels of soluble/insoluble Aβ [36]. Together with our data, revealing an early and robust loss of several nAChRs not only in the brain but also in brain mitochondria along with a proinflammatory brain state, and a potential of an α7 receptor agonist to alleviate these impairments both in acute and chronic treatment regimens, these findings document a key role of α7 nAChRs in the development of AD as well as its potential as a therapeutic target. Remarkably, AD-induced cognitive dysfunction is reduced by α7 receptor agonists at any disease stage. The underlying mechanisms are likely synaptic, including the α7-mediated activation of the Ca^2+^-dependent Ca^2+^/calmodilin-CaMKII-CREB signaling pathway, with the subsequent upregulation of the pCREB-dependent genes (e.g. c-Fos, NMDA receptors, p-TrkB, BDNF) [61], critically important for learning and memory [36, 59]. When starting, as in the current study, the α7 receptor agonist-mediated treatment during the early disease stage, additional advantages are obtained. These include a reduction of proapoptotic Cyt *c* release from the brain mitochondria, reduced levels of pro- and enhanced levels of anti-inflammatory cytokines (i.e. reduced neuroinflammation), and a reduction in the total amyloid load.

Interestingly, analyses of a mouse model of autoimmune encephalomyelitis revealed a strong anti-inflammatory effect of the PNU282987 treatment, likely caused by the inhibition of the NLRP3 inflammasome and a decreased production of IL-6, IL-1β, IL-18, and TNF-α [24, 62]. As α7 nAChRs are also expressed in astrocytes and microglia [23, 24, 53], both of which are critically involved in AD-mediated neural network hyperactivity in APPswe/PS1_G384A_ mice [12, 13], similar mechanisms might cause the reduction of proinflammatory cytokines in our study. Associations between neural network hyperactivity, high brain levels of pro-inflammatory cytokines, likely caused by Ca^2+^-dependent activation of NLRP3 inflammasome, and episodic memory impairment were also observed during an LPS-induced peripheral inflammation, and this state was accompanied by a decrease in brain and mitochondrial levels of α7 nAChRs as well as an increase of α7-bound Aβ_1-42_ [63–66], underscoring the involvement of α7 nAChRs in the inflammatory reactions within the brain. Similarly, agonist-mediated modulation of α7nAChRs was shown to prevent astrocyte activation, cytokine release, and the loss of dopaminergic neurons in the rodent models of Parkinson’s disease [53].

In conclusion, the data obtained strongly suggest that α7 nAChRs are critically important for maintaining cognitive capabilities in conditions of amyloidosis and neuroinflammation and, besides their direct interaction with Aβ, are strongly and consistently downregulated at the very onset of AD. The therapeutic use of an α7-selective agonist is well tolerated in chronic settings, profoundly upregulates the expression of all abundant brain nAChRs, reduces both free and the α7-coupled Aβ within the brain, has anti-inflammatory and antiapoptotic effects, and potently upregulates cognition.

## Data availability

The data in the present study are available from the corresponding author upon reasonable request.

## CRediT authorship contribution statement

**Olena Lykhmus**: investigation, visualization, validation, draft editing. **Wen-Yu Tzeng:** formal analysis, visualization, investigation, writing - original draft. **Lyudmyla Koval:** investigation. **Kateryna Uspenska:** investigation. **Elizabeta Zirdum**: investigation. **Olena Kalashnyk**: investigation, draft editing. **Olga Garaschuk:** conceptualization, writing – original draft, review and editing, supervision. **Maryna Skok:** conceptualization, methodology, investigation, writing – original draft, supervision.

## Acknowledgments

We thank E. Zirdum, A. Weible, K. Schmidt and B. Ott for technical assistance and Y. Kovalchuk for his help with analyses of behavior.

---

**Figure S1.** Longitudinal changes in the levels of nAChR subunits in the brain or brain mitochondria of the WT and AD mice. (**A**, **B**) Bar graphs, illustrating the age-dependent changes in the levels of different nAChR subunits in the detergent lysates of the whole brain (**A**) and brain mitochondria (**B**) of WT mice. (**C**, **D**) similar data as in (**A**, **B**) shown for AD mice (n = 4 per group, two-way ANOVA followed by Bonferroni’s multiple comparisons test, *p<0.05 for all comparisons).

**Figure S2.** The effect of 4-month-long PNU282987 treatment on the body weight and locomotor activity in the open field test. (**A**) Sample trajectory plots of AD mice after 4-month-long treatment with either PBS (right) or PNU282987 (left) during the 10-minute-long test period. (**B**-**D**) Bar graphs illustrating the distance traveled (**B**), average speed (**C**) and number of fecal boli released (**D**) during the open field test. (**E**) Graph illustrating the weekly-measured body weight during the 4-month-long treatment. Data are shown as mean ± SEM, **B**-**D**: Student’s t-test, p>0.5; **E**: two-way ANOVA, p=5.6x10^-2^; n=6 mice per group.

