## Supplementary figures and images for "Impairment of brain function in a mouse model of Alzheimer’s disease during the pre-depositing phase: the role of α7 nicotinic acetylcholine receptors"

### Supplementary Figures

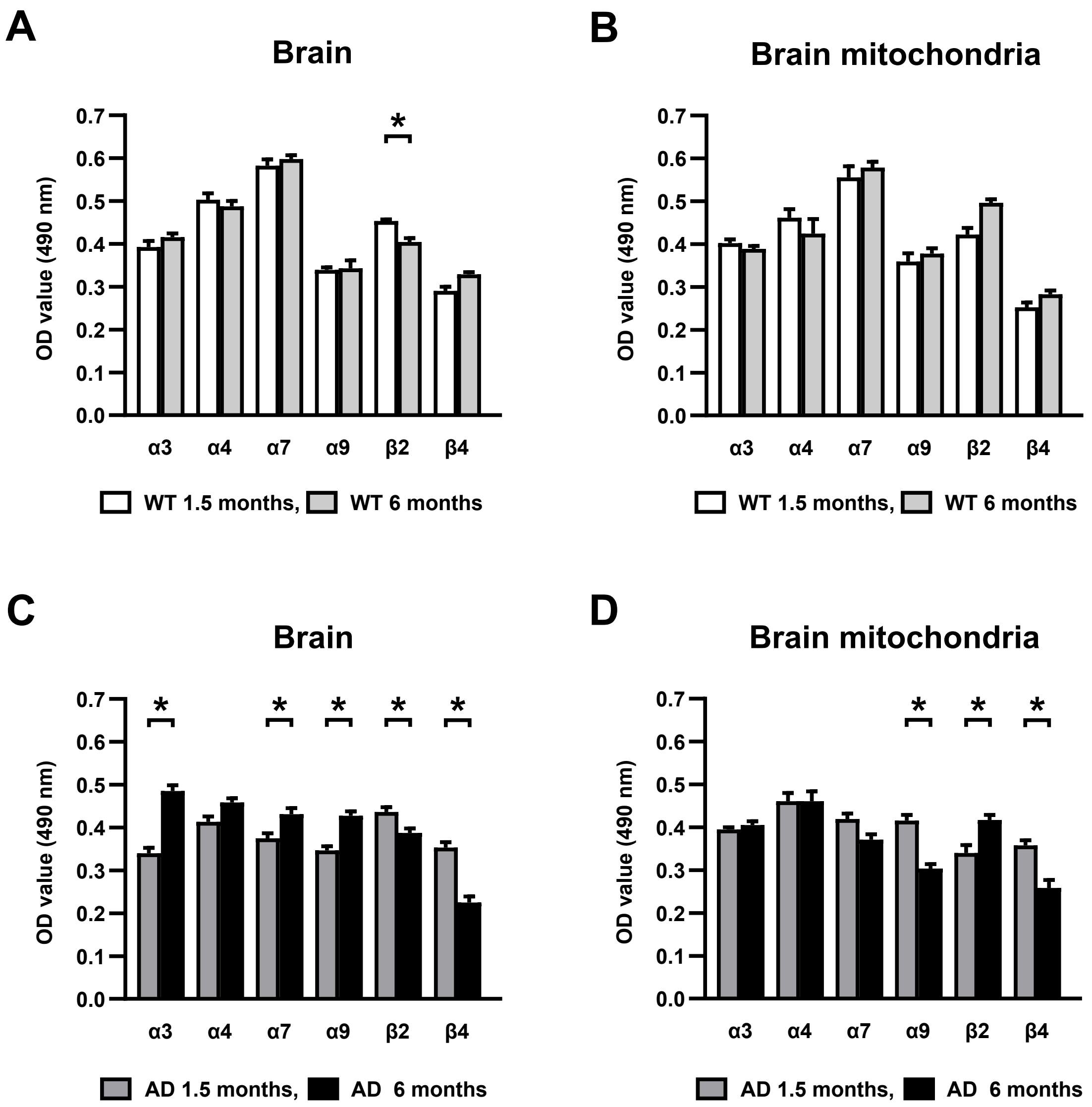

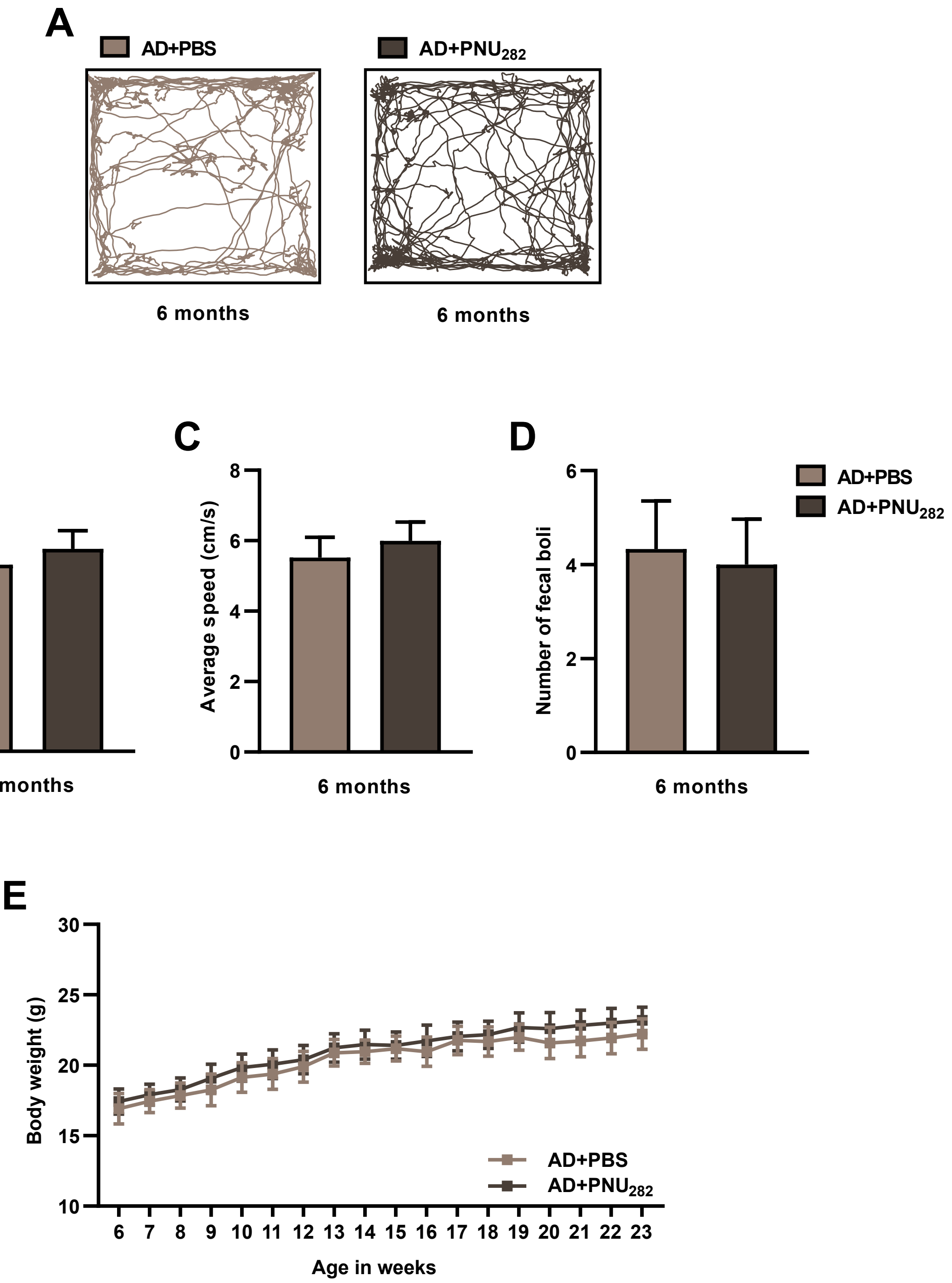
